# Combination of structural and functional connectivity explains unique variation in specific domains of cognitive function

**DOI:** 10.1101/2021.10.20.463183

**Authors:** Marta Czime Litwińczuk, Nelson Trujillo-Barreto, Nils Muhlert, Lauren Cloutman, Anna Woollams

## Abstract

The relationship between structural and functional brain networks has been characterised as complex: the two networks mirror each other and show mutual influence but they also diverge in their organisation. This work explored whether a combination of structural and functional connectivity can improve predictive models of cognitive performance. Principal Component Analysis (PCA) was first applied to cognitive data from the Human Connectome Project to identify components reflecting five cognitive domains: Executive Function, Self-regulation, Language, Encoding and Sequence Processing. A Principal Component Regression (PCR) approach was then used to fit predictive models of each cognitive domain based on structural (SC), functional (FC) or combined structural-functional (CC) connectivity. Self-regulation, Encoding and Sequence Processing were best modelled by FC, whereas Executive Function and Language were best modelled by CC. The present study demonstrates that integrating structural and functional connectivity can help predict cognitive performance, but that the added explanatory value may be (cognitive) domain-specific. Implications of these results for studies of the brain basis of cognition in health and disease are discussed.

**Highlights:**

- We assessed the relationship between cognitive domains and structural, functional and combined structural-functional connectivity.
- We found that Executive Function and Language components were best predicted by combined models of functional and structural connectivity.
- Self-regulation, Encoding and Sequence Processing were best predicted by functional connectivity alone.
- Our findings provide insight into separable contributions of functional, structural and combined connectivity to different cognitive domains.

## 1. Introduction

Cognitive neuroscience generally seeks to develop an understanding of neural substrates of cognition and adaptive behaviour. One approach to the study of the brain is to characterise it as a network of brain regions and connections between them (Fornito et al., 2016; Sporns et al., 2005). Following this approach, structural connectivity (SC) of a brain network describes the patterns and the integrity of white matter connections between neural populations (Sporns et al., 2005), whereas functional connectivity (FC) of a brain network describes patterns and strength of temporal associations of activation patterns across remote brain regions (Bullmore & Sporns, 2009; Friston, 2002). In recent years, efforts have been made to investigate how SC and FC relate to each other and how this relationship may affect cognitive function and health (Bullmore & Sporns, 2009; Rykhlevskaia et al., 2008). A key question is whether SC and FC provide complementary or, alternatively, overlapping information for explaining cognition and behaviour (de Kwaasteniet et al., 2013; Guye et al., 2010; Hahn et al., 2013; Salami et al., 2014; van den Heuvel & Fornito, 2014; Wang et al., 2016).

In general, the research that relates brain structure and function is motivated by the fact that the human brain activates and operates on the scaffold of neurons and neuronal connections. Consequently, SC and FC must be related to some degree. In support of this proposal, evidence demonstrates systematic coupling of lifespan changes in SC and FC (Baum et al., 2020; Romero-Garcia et al., 2014). Further, research has found striking similarity between white matter connectivity profiles and functionally meaningful parcellations of the cortex (Greicius et al., 2008; Johansen-Berg et al., 2004; Jung et al., 2017; Vázquez-Rodríguez et al., 2019). To elaborate, Johansen-Berg and colleagues (2004) have found that the SC profile of human medial frontal cortex shows abrupt changes at boarders of functionally meaningful regions. Further yet, evidence demonstrates that the most central nodes of functional networks are directly and strongly connected by white matter tracts (Greicius et al., 2008). There are three important features that characterise the brain’s structure-function relationship. First, spatially organised neuronal populations determine where activation can occur. Evidence demonstrates that the degree of functional activation of a cortical area is influenced by the physical properties of the region, including cortical volume, thickness, surface area and curvature (Chen et al., 2018; Tillisch et al., 2017). Second, the amount of locally exchanged of cortical activity is influenced by local connection density (Bassett & Bullmore, 2017; Bullmore & Sporns, 2012; Cammoun et al., 2014; Powell et al., 2006). Third, exchange of activities across remote regions is more efficient when it is supported by long-range white matter tracts with excellent integrity (Bullmore & Sporns, 2012; Liu et al., 2017; Taubert et al., 2011).

The most direct evidence of structural influence on neural activation comes from neurostimulation studies. It has been shown that the presence and integrity of direct white matter connections can modulate the impact of transcranial direct current stimulation on global patterns of neural function and cognitive outcomes (Li et al., 2019; Lin et al., 2017). These findings demonstrate that brain structure can directly impact the strength of FC between select regions. On the other hand, it has been hypothesised that regions that fire together, will eventually wire together through plasticity mechanisms (Draganski et al., 2004; Gaser & Schlaug, 2003; Hölzel et al., 2011). In support of this hypothesis, studies have found that repeated engagement or suppression of brain activity in specific brain regions can result in corresponding anatomical changes. For example, in healthy populations extended behavioural training (Gu & Kanai, 2014; May, 2011) and environmental stressors (Czéh et al., 2006; Ortiz & Conrad, 2018; Radley et al., 2015) tend to repeatedly engage brain activity in specific brain regions, and such prolonged activity results in corresponding cortical and structural connectivity changes. In clinical research, it has been found that therapy for developmental dyslexia can produce changes in cortical volume (Krafnick et al., 2011); and intense speech therapy for chronic stroke patients with Broca’s aphasia result with strengthening of white matter connections (Wan et al., 2014). Similarly, prolonged pharmaceutical interventions targeting brain function, can also impact the physical properties of the targeted neuronal populations. For example, in clinical settings, prolonged medication has induced functional and structural abnormality in regions that were previously functionally related to relevant healthy cognitive function (Fu et al., 2013; Thomaes et al., 2014). Perhaps the most striking illustration of the complex relationship between structure and function comes from administration of hormonal contraceptives to healthy adults. A recent systematic review demonstrates that such intervention results with changes to structural volume, to the intensity of neural activity and it results with affective and cognitive changes (Bronnick et al., 2020). Interestingly, most work has been focused on FC strength, but more recent evidence suggests that the topography of functional nodes is also important for cognition (Kong et al., 2019), which suggests that spatial constraints are also consequential for brain function. This evidence demonstrates that there is a diversity of (not mutually exclusive) potential brain mechanisms underlying these findings, and all are likely to be involved in the evolving neural instantiation of cognitive processing. This highlights the complexity of the pervasive interplay between brain structure and function and suggests a high degree of overlapping in their roles with regards to cognition.

Although there are patterns of overlap across SC and FC, substantial evidence demonstrates that each displays unique features that can be relevant to cognition and behaviour. For example, studies have demonstrated that regions that are not directly connected by white matter can show similar patterns of activity, which suggest that they are (indirectly) functionally connected (Ashourvan et al., 2019; Friston, 2002; Hagmann et al., 2008; Honey et al., 2009; Honey et al., 2010; Liao et al., 2015; Røge et al., 2017; Sun et al., 2012; Thomas et al., 2009). Additionally, parcellations of the human brain produced based on combined structural-functional information only show approximately 30% of overlap with structurally defined parcels (Keyvanfard et al., 2020). In a more recent publication, Mansour et al. (2021) have demonstrated that FC fingerprints are more predictive of cognitive function, while SC ones are more able to differentiate identity of individuals. This evidence suggests that, alongside shared information, the SC and FC show unique features which can impact cognition in distinct ways. Therefore, we need to understand the potential benefits of using an integrative approach, and discover exactly how the two networks diverge.

The ultimate goal of understanding neural structure and function is to provide an account of human thought and behaviour. Although quantitative simultaneous consideration of neural structure and function is relatively rare in the imaging literature, a few recent investigations have demonstrated that cognition is supported by an interplay between SC and FC. For example, high working memory performance has been associated with strong SC and increased competition between functional subnetworks (fronto-parietal and default mode) (Murphy et al., 2020). This illustrates that the previously proposed role of brain structure as a scaffolding for brain function can influence cognitive performance. In another study, Dhamala et al. (2021) demonstrated that the integration of SC and FC information improved the accuracy of linear models of fluid intelligence, composed of executive function, attention, picture sequence memory, working memory, and processing speed. Conversely, functional information alone best modelled crystallised intelligence composed of language. This suggests that the extent of the impact of the relationship between SC and FC on cognitive performance may depend on the cognitive domain that is being analysed. However, composite intelligence scores are theory-driven and do not necessarily reflect the underlying variance. This means that they serve as good summaries of cognitive performance but it may loose on important information about cognitive health (Barch & Ceaser, 2012; Covey et al., 2011; De Felice & Holland, 2018; Rivera-Fernández et al., 2021), Consequently, further investigations should explore how the relationship between structure and function impacts performance across individual cognitive domains.

The present research used diffusion Magnetic Resonance Imaging (dMRI), resting-state functional Magnetic Resonance Imaging (rs-fMRI) and cognitive data from the Human Connectome Project (HCP) to investigate how combining SC and FC (integrative approach) benefits the understanding of the neurobiological basis of cognitive function. To do this we carried out a model comparison analysis to arbitrate between competing (linear) predictive models of specific cognitive domains. For each cognitive domain, the competing models differed in the type of connectivity data used as predictors, namely: (i) SC, (ii) FC or (iii) Combined Connectivity (CC) which combined SC and FC. This model comparison approach allowed us to select the domain-specific best predictive model, based on connectivity measures of structure and function. In doing so, we tested the hypothesis that, including information from both brain structure and function (i.e. CC) would improve prediction of performance in a cognitive domain over and above that provided by SC or FC in isolation.

## 2. Materials and methods

### 2.1.1. Participants

Behavioural data was obtained for 682 subjects from the 1200-subject release of HCP dataset (Van Essen et al., 2013). The sample of subjects for behavioural analysis was selected by only including those subjects who have complete behavioural data and neuroimaging data (i.e. at least one T1-weighted image, resting-state fMRI and diffusion MRI). The sample consisted of 370 females and 312 males. The subjects had age ranges of 22–25 years (N□=□130), 26–30 (N□=□315), 31–35 (N□=□232), and 36-100 years (N = 5).

Neuroimaging data was obtained for 249 unrelated subjects from the 1200-subject HCP release. For consistent treatment of behavioural and neuroimaging subject selection, one subject was excluded from the neuroimaging analysis due to incomplete behavioural data. The sample consisted of 138 females and 111 males. The subjects had age ranges of 22–25 (N□=□45), 26–30 (N□=□105), 31–35 (N□=□96), and 36-100 (N = 3).

#### 2.1.2. Measures of cognition

The present study used all behavioural data from the domain of cognition (Barch et al., 2013) obtained with tasks from the Blueprint for Neuroscience Research–funded NIH Toolbox for Assessment of Neurological and Behavioral function (http://www.nihtoolbox.org) and tasks from the Penn computerized neurocognitive battery (Gur et al., 2010). The cognitive data comprised of measures of verbal and non-verbal episodic memory, cognitive flexibility, inhibition, language, fluid intelligence, processing speed, impulsivity, spatial orientation, attention and working memory. Analysed tasks include: Picture Sequence Memory, Dimensional Change Card Sort, Flanker Inhibitory Control and Attention Task, Penn Progressive Matrices, Oral Reading Recognition, Picture Vocabulary, Pattern Comparison Processing Speed, Delay Discounting, Variable Short Penn Line Orientation Test, Short Penn Continuous Performance Test, Penn Word Memory Test, and List Sorting. Supplementary material 1 presents a table summary of each tasks’ cognitive subdomain and a brief outline of its cognitive demands. The present work has used scores that were not adjusted by age, as the age range is narrow (22-35). Participant performance was normalised using the NIH Toolbox Normative Sample (18 and older). Following normalisation, a score of 100 indicates average performance and a score of 115 or 85 indicates performance 1 SD above or below the national average respectively. For Penn Progressive Matrices, Penn Word Memory Test and Variable Short Penn Line Orientation Test the median reaction time for correct responses was divided by accuracy, to obtain a measure of overall efficiency of task performance (Liesefeld & Janczyk, 2018). For the Delay Discounting task, the area under the curve was used from both $200 and $40 000 versions of the task.

#### 2.1.3. Minimally processed Neuroimaging data

The HCP provides minimally processed neuroimaging data that was used here, the data acquisition and processing pipeline has been discussed in detail by (Glasser et al., 2013). All neuroimaging data was collected with a 3T Siemens “Connectome Skyra” scanner that uses the Siemens 32-channel RF receive head coil and with SC72 gradient insert (Ugurbil et al., 2013). Here, we utilised Version 3 of the minimal processing pipeline implemented with FSL 5.0.6 (Jenkinson et al., 2012) and FreeSurfer 5.3.0-HCP (Dale et al., 1999).

T1 weighted MR images were acquired with a 3D MPRAGE sequence (TR = 2400 ms, TE = 2.14, TI = 1000 ms, flip angle = 8°, FOV = 224 by 224 mm, voxel size = 0.7 mm isotropic). Rs-fMRI data was collected using the gradient-echo EPI (TR = 720 ms, TE = 33.1 ms, flip angle = 52°, FOV = 208 by 180 mm, 70 slices, thickness = 2.0 mm, size = 2.0 mm isotropic). Scans were collected in two sessions, each lasting approximately 15 minutes. The rs-fMRI data was collected both in left-to-right and right-to-left directions. In addition, in the original data, spin echo phase reversed images were acquired for registration with T1 images and the spin echo field maps were acquired for bias field correction. Diffusion weighted MR images were acquired with spin-echo EPI sequence (TR = 5520 ms, TE = 89.5 ms, flip angle = 78°, refocusing flip angle = 160°, FOV = 210 by 180 mm, 111 slices, thickness = 1.25 mm, size = 1.25 mm isotropic). Each gradient consisted of 90 diffusion weighting directions plus 6 b=0. There were 3 diffusion weighed shells of b=1000, 2000, and 3000 s/mm^2^. SENSE1 multi-channel image reconstruction was used (Sotiropoulos et al., 2013).

### 2.2. Additional processing of Neuroimaging data

#### 2.2.1 Structural data and SC calculation

As additional steps to the minimal processing pipeline, the diffusion data was put through BEDPOSTX procedure in FSL, which runs Markov Chain Monte Carlo sampling to estimate probability distributions on diffusion parameters at each voxel. This information was used in the FDT module of FSL to run ROI-to-ROI probabilistic tractography with ProbtrackX. Tractography was run between parcels obtained with a high resolution functionally defined brain parcellation with 278 parcels (Shen et al., 2013). During tractography, 5000 streamlines were initiated from each voxel with step length of 0.5 mm (Behrens et al., 2007; Behrens et al., 2003; Jenkinson et al., 2012). Streamlines were constrained with curvature threshold of 0.2, maximum of 2000 steps per streamline and volume fraction threshold of subsidiary fiber orientations of 0.01. A SC matrix between regions was constructed by first counting the number of streamlines originating from a seed region *i* that reached a target region *j* (*M*_*ij*_). These counts are asymmetric since the count of streamlines from region *i* to *j* is not necessarily equal to the count of streamlines from region *j* to *i* (*M*_*ij*_ *≠ M*_*ji*_), but they are highly correlated for all subjects (lowest Pearson’s Correlation was 0.76, p < 0.001). Based on these counts, the weight *W*_*ij*_ (entries of the SC matrix) between any two pairs of regions *i* and *j* was defined as the ratio of the total streamline counts in both directions (*M*_*ij*_ *+ M*_*ji*_), to the maximum possible number of streamlines that can be shared between the two regions, which is *(N*_*i*_ *+ N*_*j*_) * 5000 (where *N*_*i*_ and *N*_*j*_ are the number of seed voxels in regions *i* and *j*, respectively):,

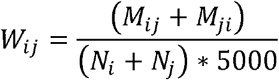

Similar to previous studies, the weight *W*_*ij*_ can be interpreted as capturing the connection density (number of streamlines per unit surface) between nodes *i* and *j*, which accounts for possible bias due to different sizes of the seed regions (Friston et al., 2008; Ingalhalikar et al., 2013). Note that the SC matrix defined based on these weights is symmetric because swapping around the regions’ indices does not change the result; and it is also normalised between 0 and 1, because the maximum value of the numerator can only be reached when all streamlines originating from each of region reach the other region, so that *M*_*ij*_ *= N*_*i*_ * 5000 and *M*_*ji*_ *= N*_*j*_ * 5000, which gives *W*_*ij*_ = 1. Evidence suggests that structural connectivity is most sensitive to individual differences with moderate-to-high thresholding (Buchanan et al., 2020) and produces least false positive and negative results (de Reus & van den Heuvel, 2013), therefore an 80% proportional threshold was applied. Supplementary material 2 presents the results of analysis conducted on dense connectivity.

#### 2.2.2. Functional data and FC calculation

The minimally processed images were obtained for rs-fMRI to compute functional connectivity based on pair-wise correlations (Glasser et al., 2013). Next, the following steps were taken to further process data using the CONN Toolbox (Whitfield-Gabrieli & Nieto-Castanon, 2012) with use of the standard functional connectivity processing pipeline (Nieto-Castanon, 2020). Briefly, the tissues were segmented into separate parcels, corresponding to grey matter, white matter, cerebrospinal fluid and non-brain tissues. Outlier detection of functional images for scrubbing was performed with Artifact Detection Tools (ART, https://www.nitrc.org/projects/artifact_detect/). Images were smoothed with a 6 mm Full Width at Half Maximum Gaussian kernel. The data was then high pass filtered (0.01 Hz cut-off), and a low pass filtered (0.10 Hz cut-off). Finally, an anatomical component-based noise correction procedure (Behzadi et al., 2007) was conducted to remove the following artefactual components from the data: noise components from cerebral white matter and cerebrospinal areas, subject-motion parameters (Friston et al., 1996), identified outlier scans (Power et al., 2014), and constant and first-order linear session effects (Whitfield-Gabrieli & Nieto-Castanon, 2012). FC analysis was performed based on the same high resolution brain parcellation used in the SC computations (Shen et al., 2013). The average blood oxygenation level-dependent signal in each ROI was obtained and the pairwise (ROI-to-ROI) correlation of the averaged signals was calculated. Since the CONN toolbox produces Fisher’s Z-scores (Fisher, 1915), a hyperbolic tangent function was used to reverse the Fisher’s transformation, and obtain original correlation values ranging between −1 and 1. Negative correlations were transformed to positive by taking their absolute values and a proportional 80% FC threshold was then applied (Garrison et al., 2015; van den Heuvel et al., 2017). This procedure has been shown to produce more reliable components resulting from Principal Component Analysis (PCA) of functional connectivity (Hong et al., 2020). Supplementary material 2 presents the results of analysis conducted on dense connectivity and supplementary material 3 presents the results of preserving sign of negative correlations.

## 3. Data analysis

### 3.1. Behavioural data analysis

The HCP provides a wealth of behavioural data that assess a rich variety of cognitive tasks. Consequently, it is possible that different behavioural measures draw on shared cognitive processes. Principal Component Analysis (PCA) to was used estimate orthogonal rotated components (RC) each reflecting specific latent cognitive domains (Butler et al., 2014; Hoogendam et al., 2014; Levin et al., 2013; Schumacher et al., 2019). This PCA-based cognitive domain extraction was carried out on the cognitive data of 433 out of the total of 682 participants, by setting aside the 249 participants (for which the Neuroimaging data was available) used later for constructing the different predictive models of cognition. Specifically, after standardising the data (z-score) the most stable number of PCA components with eigenvalues equal to or greater than 1 (Guttman-Kaiser rule) was estimated by randomly selecting 90% of the 433 participants for the PCA estimation and repeating the procedure 10000 times. An eigenvalue of 1 has been proposed as a reasonable lower bound for selecting PCA components in psychology research (usually leading to 80% of explained variance and interpretable components) and has been shown to have some optimal reliability properties, as being a necessary and sufficient condition for a principal component to have positive Kuder-Richardson reliability (Kaiser, 1960). The complementary stability analysis described above showed 5 to be the most stable number of components (occurring in 78.91% of cases), explaining 63.831% of the variance on average (standard deviation 3.45%). The estimated RCs were then used to compute RC scores of the remaining 249 subjects, which were taken as response variables for the subsequent model construction (see next section). Since the proposed model construction approach was embedded in a k-fold cross-validation (CV) setting, splitting the cognitive data for cognitive component estimation as above, prevents data-leakage during the CV procedure. This is because the estimation of the cognitive components on the separate sample of 433 subjects, effectively makes the subsequent computation of the component scores of the remaining 249 subjects, independent of the CV procedure.

### 3.2. Model Construction and Model Comparison

#### 3.2.1. Overall approach

Model construction was based on a cross-validated (CV) Principal Component Regression (PCR) (CV-PCR) approach, where separate linear predictive models of individual cognitive domains (5 domains), were fit using either SC, FC or CC as predictors. The full CV-PCR was run for every combination of predictor type (SC, FC or CC) and cognitive domain response variable (each cognitive RC score). This resulted in a total of 15 models to be estimated (5 cognitive domains * 3 types of connectivity predictors). Finally, the model comparison analysis between the three types of connectivity models obtained for each cognitive construct was based on the Akaike Information Criterion (AIC) (Akaike, 1974). The AIC balances the goodness-of-fit of the model (model accuracy) against its complexity (the number of parameters involved in the model), so that reduced AIC is associated with improved model quality.

#### 3.2.2. Principal Component Regression

The PCR method solves a regression problem of a response variable (here a cognitive RC score associated with a given construct) on multiple explanatory or predictor variables (here SC, FC or CC values) in three steps:

##### Step 1

the PCA of predictor variables (connectivity information) is carried out to deal with multi-collinearity of predictors and to reduce the dimensionality of the problem by selecting the orthogonal principal components (PC) explaining 80% of cumulative variance of the predictors;

##### Step 2

the selected PC scores are used as new predictors to solve the regression problem in the reduced PCA space;

##### Step 3

the estimated regression coefficients in PCA space are projected back to the original connectivity space using the corresponding PC weights, to obtain one regression coefficient for each connection (network edge).

The SC and FC predictor matrices were defined by first vectorising the elements of the lower triangle of the respective connectivity matrices of each subject to obtain one row connectivity vector per subject for each connectivity type. The connectivity vectors of all subjects, were then stacked row-wise for each connectivity type to obtain separate SC or FC predictor matrices, each of dimension number of subjects (subjects’ dimension) by number of connection edges (connectivity dimension). The predictor matrix for the CC case was constructed by concatenated the SC and FC matrices column-wise. This means that in the case of the CC model, the PCA in step (i) of the PCR was applied to the concatenated SC and FC data across the connectivity dimension. During step (i) 80% of predictor variance was explained by 94 SC components, 123 FC components, and 121 CC components.

One possible limitation of PCR is that the PCA in step 1 favours components to be selected based on explained connectivity variance across subjects, even if those components don’t have high predictive value. To address this, in the present work the regression problem in step 2 was solved using step-wise regression (SWR), with AIC as criterion for including or excluding a component in the regression (Figure 1). This effectively amounts to a variable selection step, where a component is considered if its inclusion improves (i.e. reduces) the AIC of the model by at least 2 units. Models with AIC difference less than 2 units are considered to be statistically equivalent (Burnham & Anderson, 2002, p. 70). Therefore, by using this cut-off value, we effectively implement an automatic Occam’s Razor; if the increase in complexity (by adding a component) improves AIC, but results in a statistically equivalent model (AIC reduction less than 2) then the simpler model (i.e. without adding the component) is preferred.

**Figure 1.**
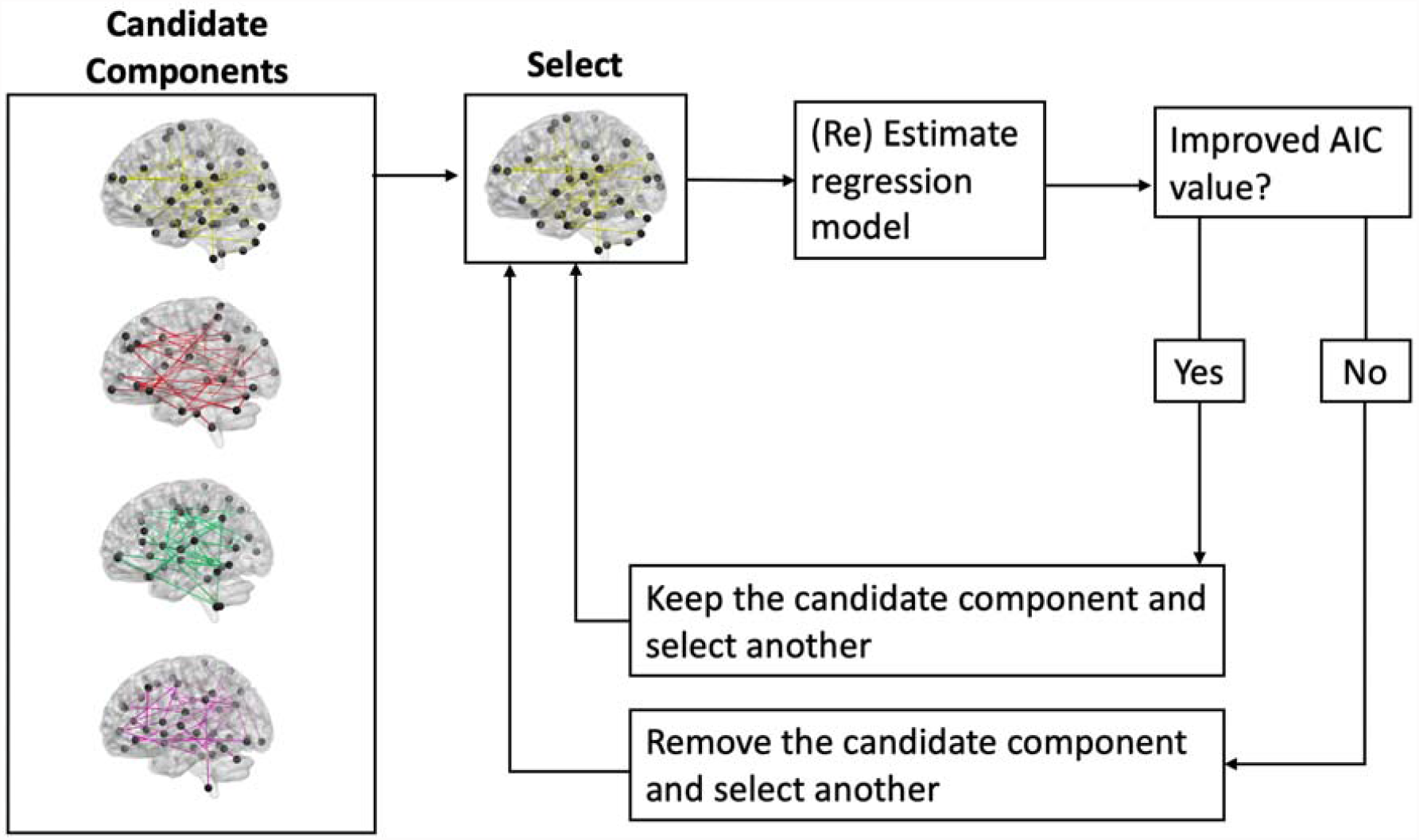
A visual illustration of the stepwise regression method applied to the connectivity components. The process is repeated until AIC value cannot be improved (reduced) any further and all candidate components have been considered by the model.

#### 3.2.3. Cross validation approach

To address overfitting and to assess the generalisation power of the models, the PCR was embedded in a randomised repeated (500 times) 10-fold CV procedure using 200 participants. The remaining 49 participants were used as a holdout data for final model skill evaluation. (Figure 2). Specifically, in each CV split 180 participants (out of the 200) were used to train the PCR model (training fold); and 20 participants were used to evaluate the skill of the trained model on unseen data (test fold). Prior to training and testing in each CV split, the mean and standard deviation of the training fold were calculated and used to standardise (centre and scale) both the training and the test folds. This procedure ensures a consistent test step because the same operations are applied to both training and test folds, and results in standardised estimated regression coefficients. The model skill on unseen data (i.e. the CV error) was then evaluated using the regression coefficients of the trained PCR model to predict the standardised test fold, and computing the root mean square error (RMSE) of the prediction. The model structure (i.e. the PCs selected in PCR) was treated as a hyperparameter so that the optimal model was the one minimising the RMSE across all CV splits.

**Figure 2.**
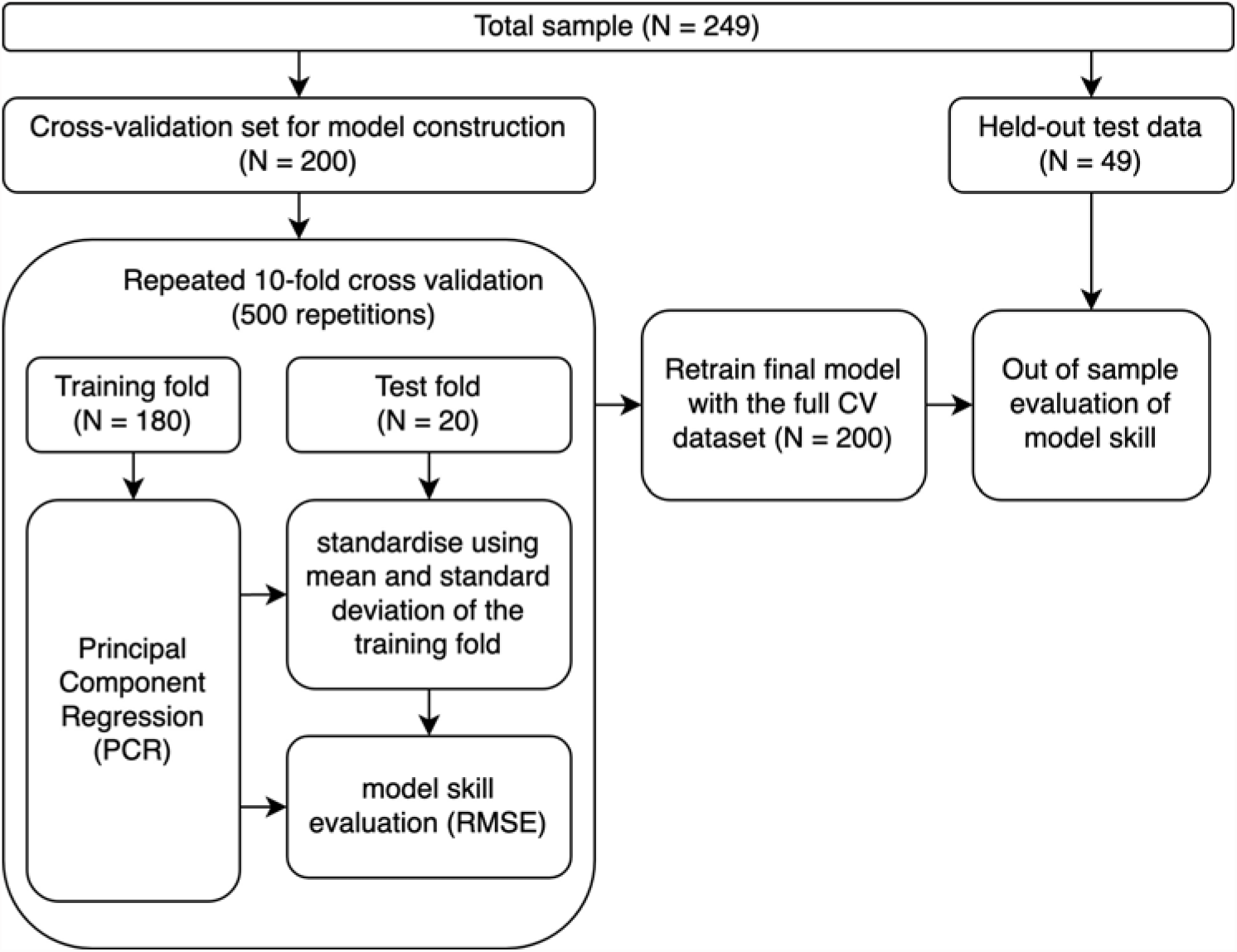
A visual illustration of the model construction and model generalisability testing pipeline. This pipeline was separately applied to each cognitive component to produce and select SC, FC or CC components and their regression coefficients. The winning model is the one that produces least prediction error, estimated with RMSE.

The whole CV data set of 200 participants was then used to re-estimate the coefficients of the optimal model. This was done by running the PCR one more time, but using the selected PC weights from the 10-fold CV (i.e. the PCs) to obtain PC scores of the full CV data set and taking them as predictors to fit standard regression (not SWR). The skill of the final model on unseen data was evaluated using the obtained regression coefficients to predict the held-out data set of 45 participants and computing the prediction error (RMSE).

#### 3.2.4. Model comparison approach

The resultant SC, FC and CC models for each cognitive domain were finally compared using AIC-based model comparison with the CC as the reference model. That is, the AIC value of the CC model was first subtracted from the AIC of the other models. Results were then interpreted so that, given any two models *M*_l_ and *M*_2_, a positive difference (Δ*AIC = AIC(M*_l_*) − AIC(M*2)) is interpreted as weak (barely worth a mention) (2-4 units), considerable (4-7 units) or strong (7-10+ units) evidence in favour of *M*_2_ (Arnold, 2010; Burnham & Anderson, 2002, p. 70). To complement this analysis, models were further assessed in terms of their explained variance in the sample of 200 participants (Ordinary R-squared)(Poldrack et al., 2020). Finally, out of sample testing was conducted on left out dataset of 49 subjects. Prediction accuracy was quantified as RMSE. All the analyses were done in MatLab 2018a (The MathWorks, Inc.).

### 3.3. Interpretation and visualisation

A specific regression coefficient in connectivity space is interpreted as the partial regression coefficient associated with a specific connectivity predictor (connection edge). Therefore, in order to identify the connections that are significantly related to a cognitive domain, we tested the hypothesis of no (linear) association between each connectivity predictor and the cognitive response variable. This was done by implementing a raw data permutation (randomisation) test (10000 permutations) for partial regression coefficients (Anderson & Legendre, 1999; Manly, 2018). To account for multiple comparisons, the above permutation test was based on the “max statistic” method for adjusting the p-values of each variable (Groppe et al., 2011). Those connections for which the null hypothesis was rejected (corrected p-value ≤ 0.05) were visualised with BrainNet Viewer (http://www.nitrc.org/projects/bnv/) (Xia et al., 2013). For illustrative purposes, node sizes were adjusted to reflect node degree, calculated using the Brain Connectivity Toolbox (https://brain-connectivity-toolbox.net) (Rubinov & Sporns, 2010).

## 4. Results

### 4.1. Cognitive components underlying behavioural data

The PCA of cognitive measures yielded a 5-component solution, which explained 62% of variance in the data. The rotated component solution was used to interpret the cognitive domains reflected by each component (Figure 3). The first component was primarily composed of Dimensional Change Card Sort, Flanker and Pattern Processing Speed tasks. The component also included a moderate negative loading from Variable Short Penn Line Orientation Test. Dimensional Change Card Sort and Flanker tasks dominated the solution with strongest loadings. Since these tests were designed to measure executive function the component will be referred to as such. The second component saw unique loading from Delay Discounting tasks, which measure impulsivity and self-control ability. Therefore, this component will be referred to as Self-regulation. The third component was dominated by Oral Reading Recognition and Picture Vocabulary tasks, as well as Penn’s Progressive Matrices. Penn’s Progressive Matrices reflect fluid intelligence rather than language ability. They are different domains supported by different neural systems (Woolgar et al., 2018), but both cognitive domains have been related in childhood development (De Stasio et al., 2014; Friesen et al., 2021; Gamaroff, 2012) and education (Kaufman et al., 2009). As the result of the dominance of language tasks on this component, we will now refer to this component as Language. The fourth component was made up of Short Penn Continuous Performance and Penn Word Memory tests, which both require effective encoding of information, therefore this component was called the ‘Encoding’ component (Cabeza et al., 2008; Ciaramelli et al., 2008). Finally, the fifth component had highest loadings associated with Picture Sequence Memory and List Sorting, both of which involve processing and reconstruction of sequences. Hence, this component will be referred to as Sequence Processing.

**Figure 3.**
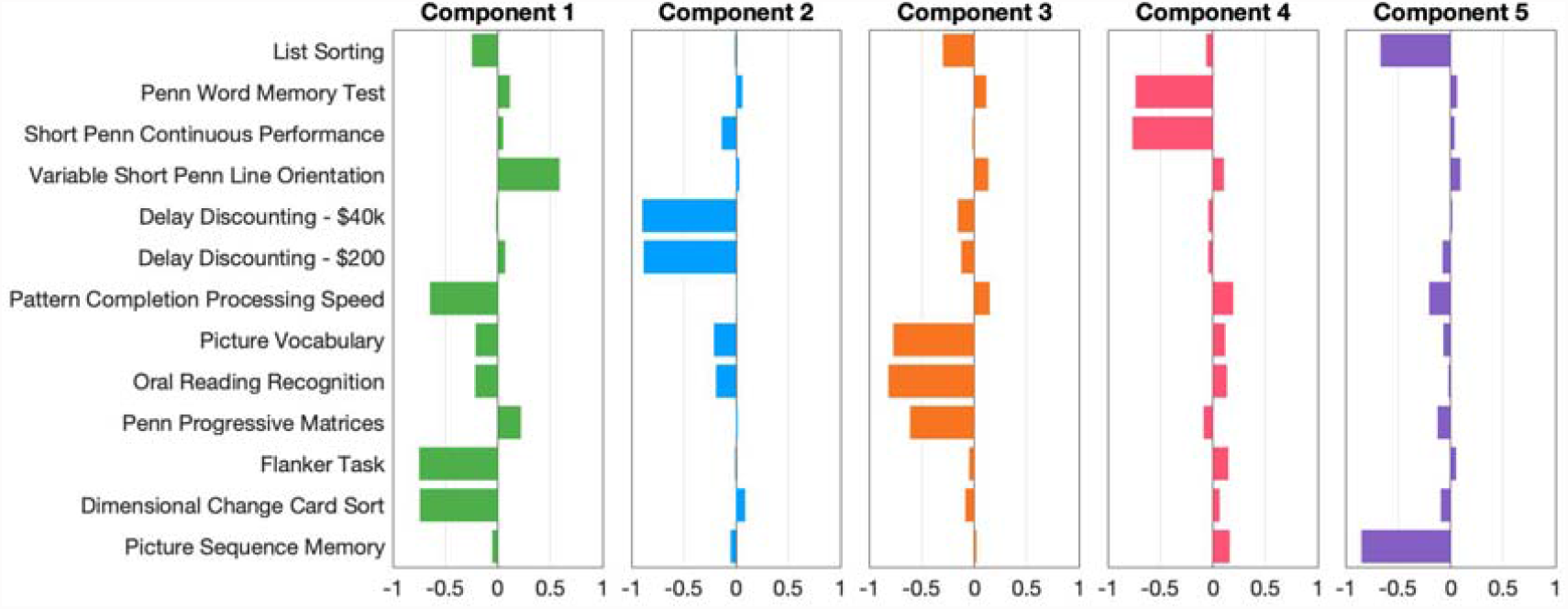
Rotated Principal Component loadings illustrated in the form of a bar graph.

### 4.2. Connectivity predictive models of cognition

#### 4.2.1. Model comparison

Across all cognitive domains there was considerable to strong preference in favour of FC relative to SC models (Executive function ΔAIC = 3.18, Self-regulation ΔAIC = 59.04, Language ΔAIC = 7.79, Encoding ΔAIC = 25.72, Sequence Processing ΔAIC = 16.26). AIC differences of SC and FC models with respect to CC model for all cognitive domains are illustrated in Figure 4. There was considerable preference in favour of CC model relative to SC model of Sequence Processing and strong preference in favour of CC models for all other cognitive domains. Compared to FC models, there was considerable empirical support of CC model of Executive Function, and strong support of CC model of Language. Conversely, there was weak support of FC model relative to CC model of Self-regulation, and strong support favouring FC models of Encoding and Sequence Processing.

**Figure 4.**
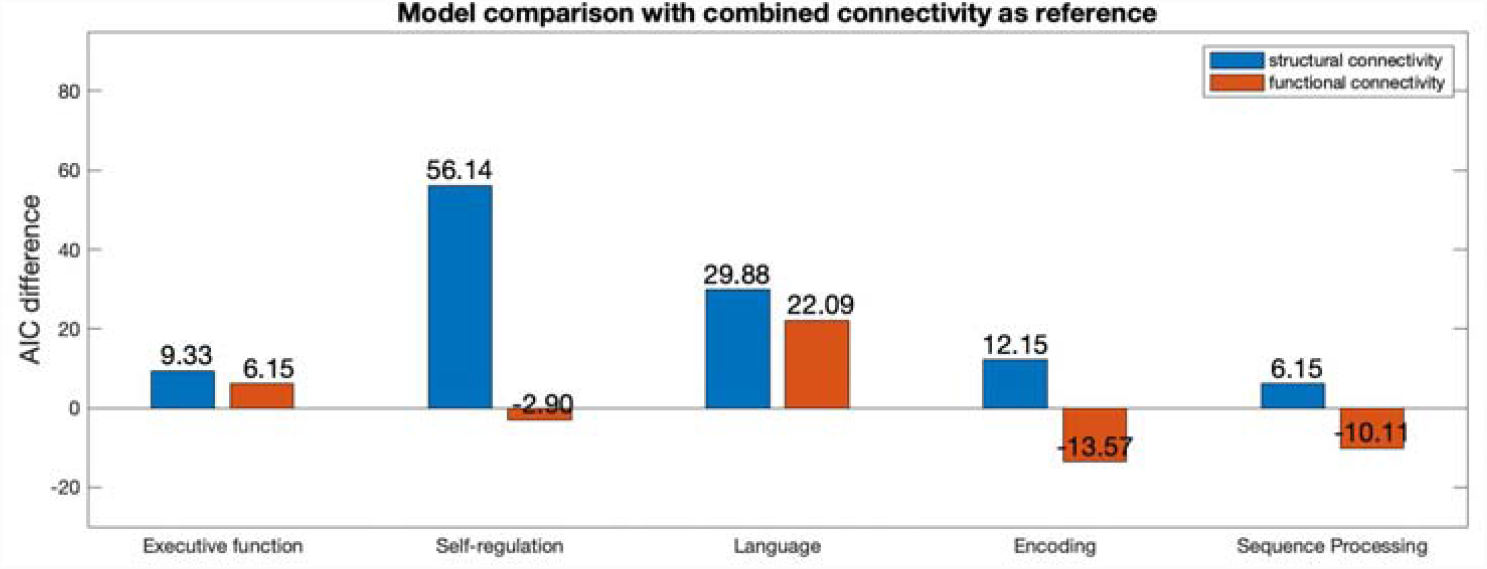
AIC-based model comparison for each cognitive domain. The CC model was used as the reference model for model comparison. Bars with positive values indicate preference for the reference model.

#### 4.2.2. Explained variance

Figure 5 illustrates the explained variance of the models under consideration. Across domains CC models achieved weak to moderate accuracy in prediction of cognitive data (Ordinary R-Squared range 0.17-0.48). The FC model explained most variance in Executive Function, Self-regulation Encoding and Sequence Processing, whereas Self-regulation and Language was best explained by the CC model.

**Figure 5.**
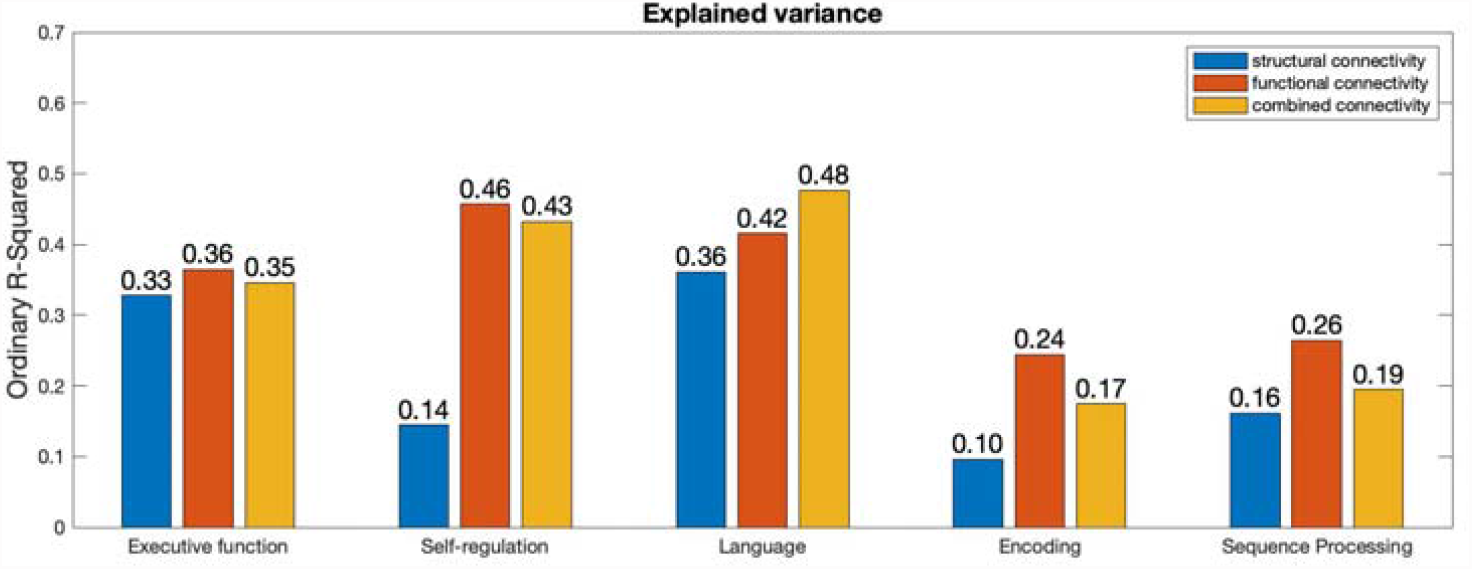
Bar graphs illustrate the variance explained by SC, FC and CC models of each cognitive domain.

#### 4.2.3. Model skill evaluation on the held-out data

The models were tested on a held-out sample of 49 participants to yield estimates of model generalisability. As illustrated in Figure 6, SC had highest generalisation skills for Executive Function, FC performed best for Self-regulation, Language, Encoding, and Sequence Processing.

**Figure 6.**
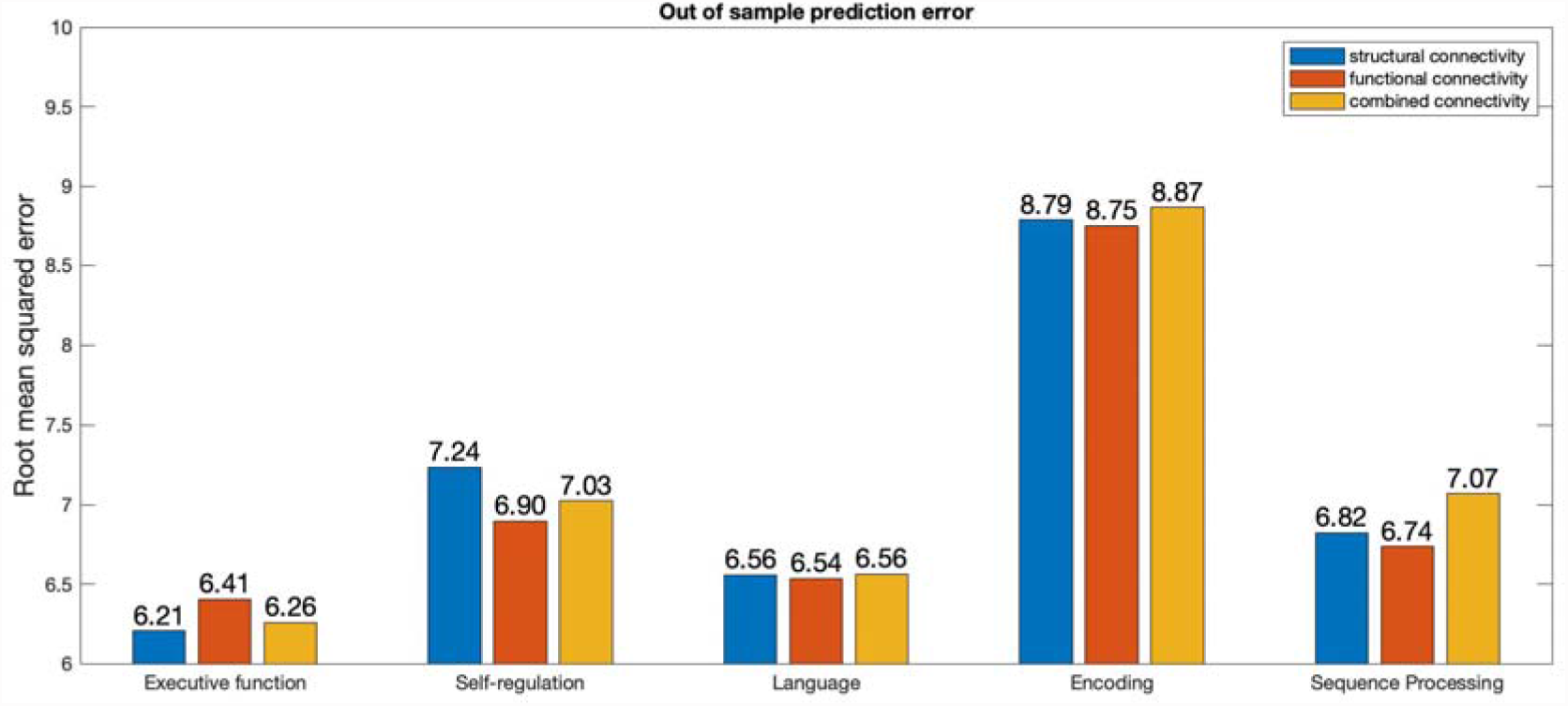
Out of sample (N = 49) prediction error, as measured by root mean squared error.

#### 4.2.4. Estimated models in connectome space

Across all cognitive domains, visualisation of the regression models in a glass brain yields a rich network of connections (Figures 7 to 11). For unimodal results, FC yields more connections associated with cognition than SC. In addition, there was no overlap between SC and FC connections supporting cognitive constructs. Compared to unimodal models, CC models yielded more structural and functional connections associated with cognition. In CC models, more functional connections were associated with cognition than structural. At most two connections overlapped across structural and functional parts of the CC models of Executive Function, Self-regulation, Language, and Sequence Processing.

**Figure 7.**
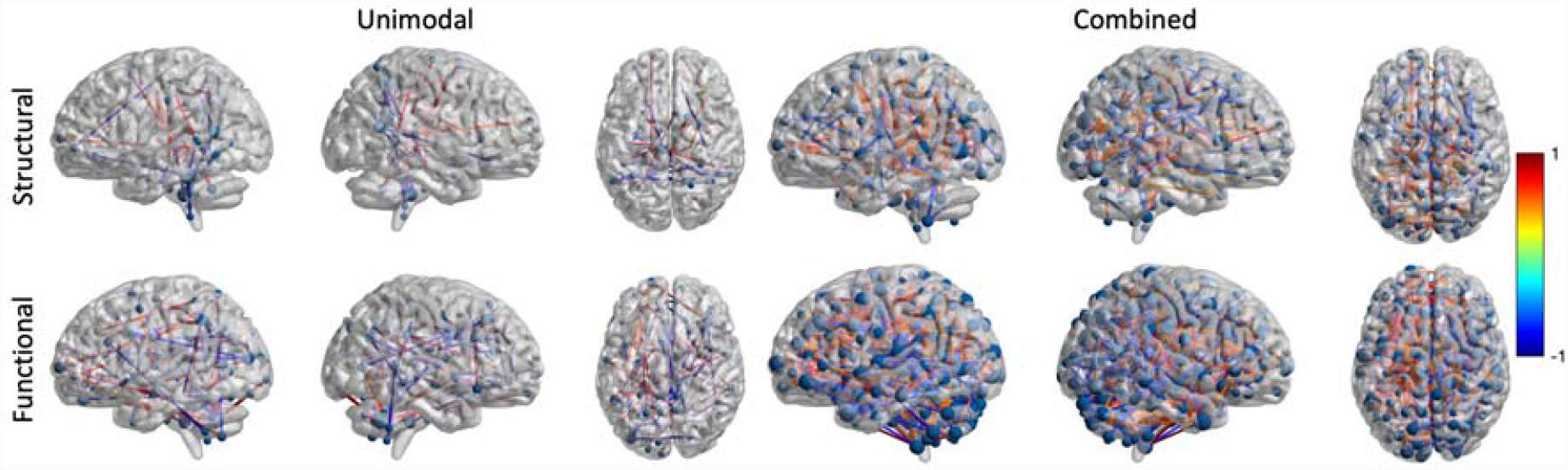
Regression models of Executive Function in connectome space. Left column illustrates unimodal models, right column illustrates combined models. In each column, top row illustrates structural connections and bottom row illustrates functional connections. The connection value range has been scaled between −1 and 1 prior to p-value based masking (p-value =< 0.05).

**Figure 8.**
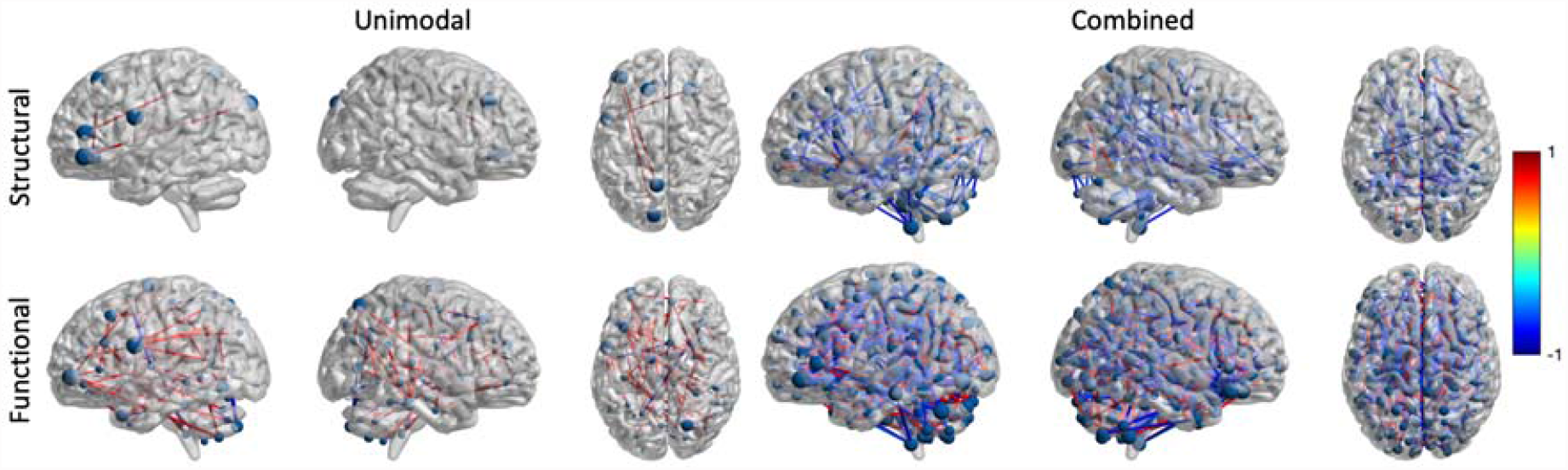
Regression models of Self-regulation in connectome space. Left column illustrates unimodal models, right column illustrates combined models. In each column, top row illustrates structural connections and bottom row illustrates functional connections. The connection value range has been scaled between −1 and 1 prior to p-value based masking (p-value =< 0.05).

**Figure 9.**
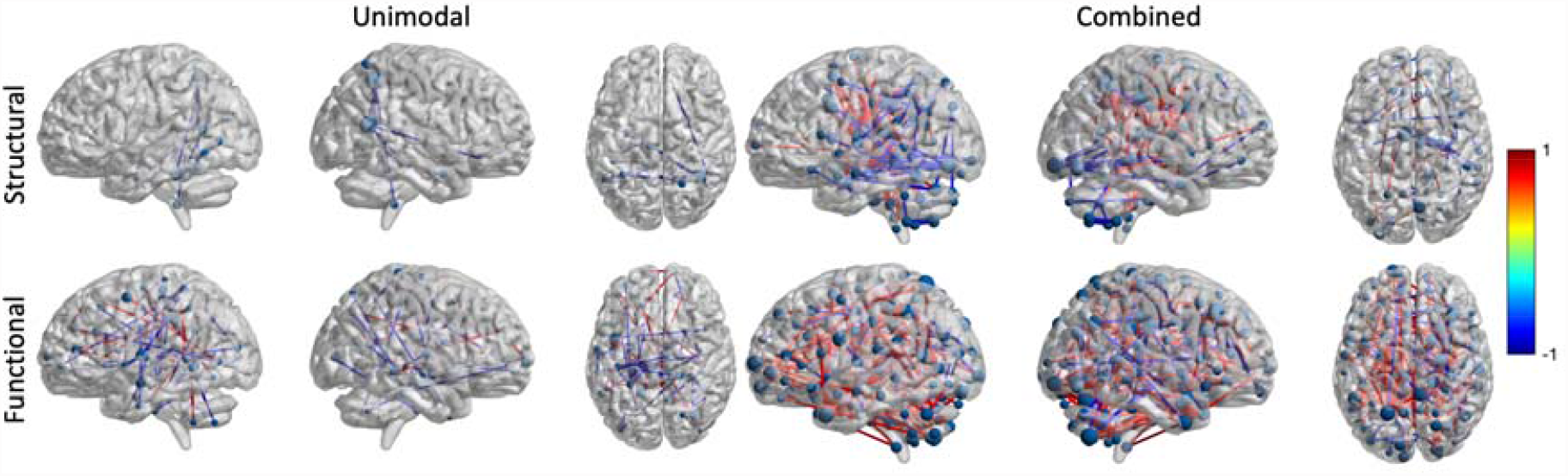
Regression models of Language in connectome space. Left column illustrates unimodal models, right column illustrates combined models. In each column, top row illustrates structural connections and bottom row illustrates functional connections. The connection value range has been scaled between −1 and 1 prior to p-value based masking (p-value =< 0.05).

**Figure 10.**
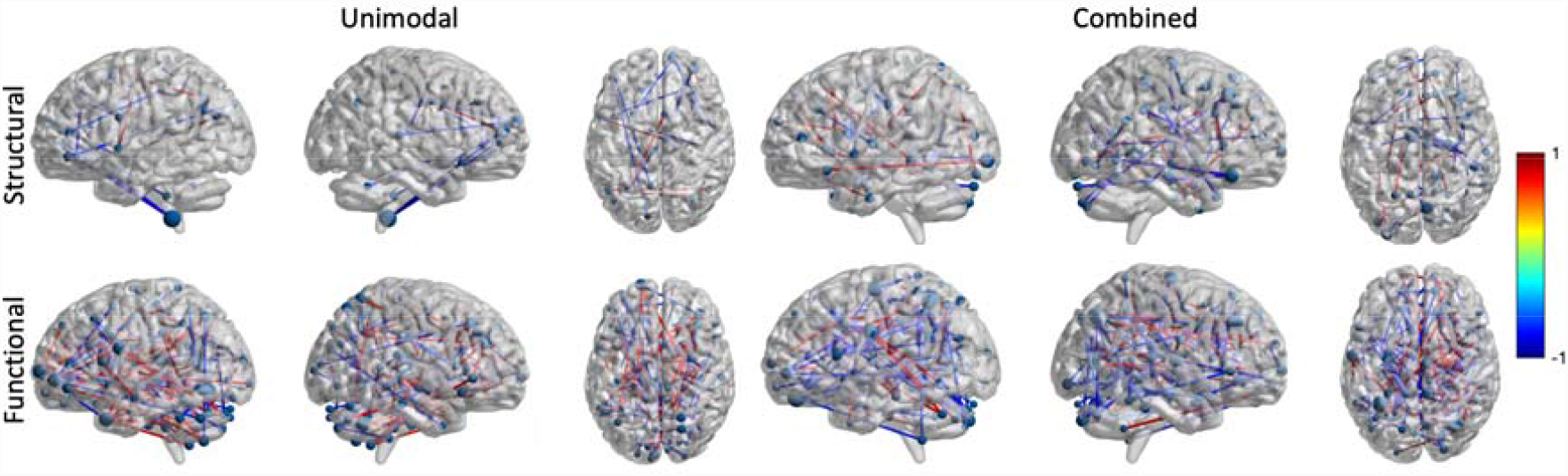
Regression models of Encoding in connectome space. Left column illustrates unimodal models, right column illustrates combined models. In each column, top row illustrates structural connections and bottom row illustrates functional connections. The connection value range has been scaled between −1 and 1 prior to p-value based masking (p-value =< 0.05).

**Figure 11.**
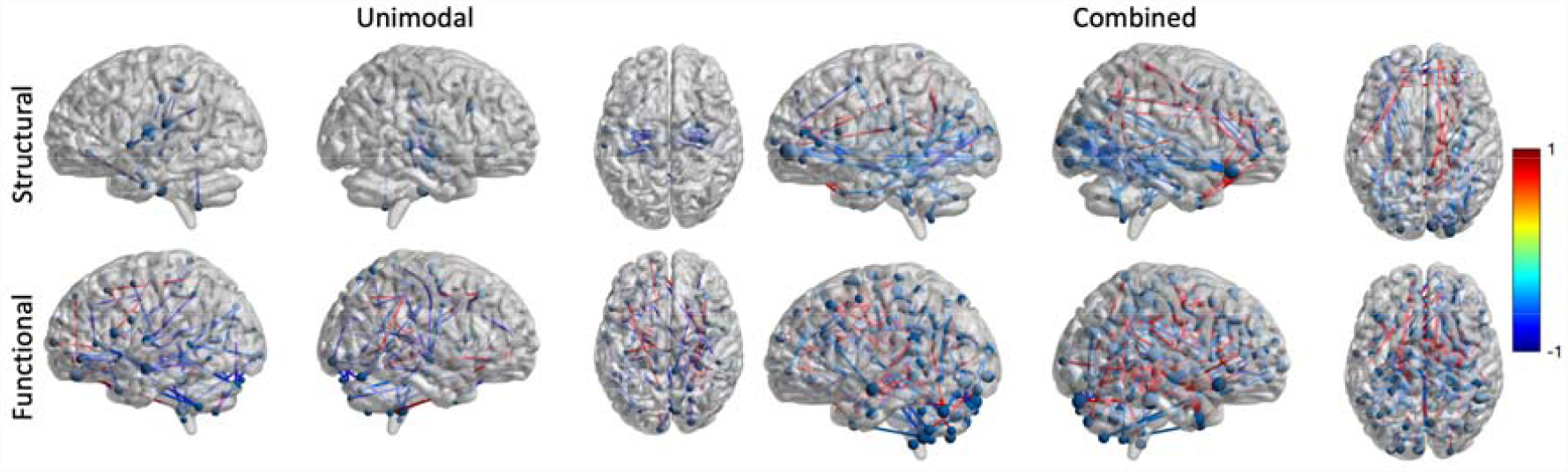
Regression models of Sequence Processing in connectome space. Left column illustrates unimodal models, right column illustrates combined models. In each column, top row illustrates structural connections and bottom row illustrates functional connections. The connection value range has been scaled between −1 and 1 prior to p-value based masking (p-value =< 0.05).

## 5. Discussion

In this study we examined whether cognitive performance was best predicted by models based on brain SC, FC or CC (i.e. combined SC and FC). First, we derived five distinct cognitive components from the HCP cognitive dataset: Executive Function, Self-regulation, Language, Encoding and Sequence Processing. The performance of SC, FC, and CC models depended on the cognitive domain. Examination of model’s AIC demonstrated that SC models of cognition were outperformed by both FC and CC models, replicating the findings from Dhamala et al. (2021) and Mansour et al. (2021). Next, CC outperformed FC at modelling of Executive Function and Language, but not Self-regulation, Encoding or Sequence Processing. Every cognitive domain was supported by a pattern of connections that was unique to structural and functional networks, replicating the findings of Zimmermann et al. (2018) and Dhamala et al. (2021). When models were used to predict unseen data, SC showed the best generalisation skills for domain of Executive Function by producing least prediction error in the unseen sample, whereas FC showed best generalisation skills for the remaining cognitive domains. The following sections first discuss the cognitive PCA analysis, and then the relative performance of each regression model.

### 5.1. Cognitive domains

In order to develop an effective understanding of the brain bases of cognition we need to quantify the relevant abilities through optimising the combination of multiple behavioural measures. Previously, researchers have combined functional and structural brain measures with summary measures of intelligence from the NIH toolbox measures, including crystallised, fluid and total intelligence, as well as ‘early’ cognition (Dhamala et al., 2021; Dubois et al., 2018; Prabhakaran et al., 1997; Robinson et al., 2021; Seidlitz et al., 2018; van den Heuvel et al., 2009; Zimmermann et al., 2018). However, these summary measures are combined based on theoretical principles, not based on shared co-variance. Consequently, they risk masking relevant variation in performance across the relevant domains.

In contrast, approaches that explore individual cognitive domains have great potential to improve our understanding and prediction of cognitive health and pathology (Barch & Ceaser, 2012; Covey et al., 2011; Rivera-Fernández et al., 2021), and have been effectively applied to predict performance in various populations, including dementia (Hackett et al., 2018; Rivera-Fernández et al., 2021) stroke aphasia (Butler et al., 2014; Mirman et al., 2019) and healthy aging (Reddy et al., 2015). We therefore considered a wide range of behavioural measures from the HCP dataset, and PCA analysis revealed five components underpinning performance on these cognitive tasks: Executive Function, Self-regulation, Language, Encoding and Sequence Processing.

### 5.2. Models of cognition

It has been proposed that every neuroimaging modality measures specific biological properties and in doing so it presents some unique information about the characteristics of the brain (Eickhoff et al., 2018). Consequently, joining information across modalities should result in a more complete representation of the state of the system. Here, we explored if combination of SC and FC benefits our understanding of performance on specific cognitive domains. To this end, we constructed regression models of cognitive performance with SC, FC and CC. To compare model quality, we assessed each model’s AIC value, accuracy (coefficient of determination) and out of sample prediction error.

When AIC values were compared, FC and CC models of cognitive performance were favoured above SC models. This finding replicates findings from previous studies that produced regression models of composite intelligence scores using SC and FC (Dhamala et al., 2021; Mansour et al., 2021). Further, we also found that fewer structural connections were associated with cognition than functional connections, which also replicates findings from Zimmermann et al. (2018). One explanation of these findings is that SC is generally sparser than FC, because FC includes indirect connectivity, observed when two remote brain regions are functionally connected without direct structural connections (Deligianni et al., 2011; Honey et al., 2009; Røge et al., 2017). Thus, SC generally presents less information about the state of the system. To explore this explanation, Zimmermann and colleagues (2018) have repeated their analysis with correction for connectome sparsity and still found that fewer structural connections supported cognition. In the present analysis, fewer components were needed to account for 80% of variance in SC than FC. This finding provides further support to the argument presented by Zimmermann et al. (2018), that there is lower variance within SC across subjects than within FC and therefore SC is less effective at exploring individual differences in cognition. However, SC is still important for understanding of cognition; AIC shows evidence in favour of FC was considerable but not overwhelming for regression models of Executive Function and Language. This suggests that specific cognitive domains may be explained well by SC. In addition, we have found that unique structural connections support cognitive performance regardless, replicating the findings of Zimmermann et al. (2018) and Dhamala et al. (2021). This pattern was observed for every cognitive domain, which suggests that unique features of SC may support cognitive performance. Taken together, these results suggest that both SC and FC are important for understanding cognitive performance.

Next, the comparison of AIC values demonstrated that Self-regulation, Encoding and Sequence Processing were best modelled by FC, whereas Executive Function and Language were best modelled by CC. The finding that joint structural-functional connectivity provides unique contribution to models of specific cognitive domains replicates previous findings from Dhamala et al. (2021) and Robinson et al. (2021). More specifically, Robinson et al. (2021) have found that the more abstract cognitive domains were best modelled by combined features of the brain. Their finding could be explained by hierarchical cortical organisation of behavioural tasks, as demonstrated by Taylor et al. (2015). Taylor and colleagues have demonstrated that concrete cognitive tasks require engagement of sensory connections. However, deeper and more distant connections are also engaged as tasks become more complex and abstract. In reference to this finding, Robinson et al. (2021) have argued that due to hierarchical cortical organisation of behavioural tasks there is a benefit to combining information about different features of the brain. Similarly, we found that highly abstract cognitive domains of Executive Function and Language were best modelled by CC. Meanwhile Encoding and Sequence Processing require more concrete memory processes. Consequently, we found that these domains were best modelled by FC. Similarly, the Delay Discounting task that loaded with Self-regulation measures person’s ability to suppress impulses and respond appropriately to the given context. Arguably, it requires some reasoning and imagination of future outcomes, which may be the reason why we find that the difference between FC and CC models is only considerable but not strong for Self-regulation.

Furthermore, we propose that the abstract domains are especially dependent on the relationship between structural and functional inter-regional connections. Taylor and colleagues (2015) have demonstrated that more abstract tasks require sensory inputs but their processing requires additional engagement of cortical connections that progress deeper into the cortex. With this finding in mind, we propose that highly abstract cognitive domains benefited from combined SC and FC because any disruption to the structural integrity of inter-regional connections impacts the speed and efficiency of signal transmission. Consequently, deeper cortical connections would accentuate the impact of structural features on highly abstract cognitive domains. This effectively explains that those domains that require deeper cortical processing are better modelled by CC because the strength of structural connections impacts the strength of functional connections.

In addition, as a further and complementary explanation of our domain-specific findings, we propose that unique features of FC influence the regression model preference. In general, connectivity PCA analysis seeks to describe the directions of greatest variance. Consequently, if joined structural-functional variance is greater than functional variance, then the joint variance will dominate the PCA solution. One source of unique functional variance is indirect FC (Deligianni et al., 2011; Honey et al., 2009; Røge et al., 2017), which can be weighed down in combined PCA if its variance is small, as compared to joint variance between structure and function. In case of Executive Function and Language, it is likely that the direction of variance of indirect connectivity is weaker than the joined structural-functional variance in the CC models.

Our work faces some limitations that must be addressed by future research. Here for the first time, PCA was applied to structural and functional connectivity as a dimension reduction method, referred to by us as combined PCA. Components of this analysis reflect common and unique directions of variance between brain structure and function. It is not clear from the present analysis whether components reflect unique information reflecting the interaction between brain structure and function or whether they reflect a weighted average of the two modalities. Future work will have to explicitly address how the two modalities interact and what contributions cross-modal interactions make to models of cognition. However, the joint PCA approach presented here has produced novel insights by establish that information concerning the relationship between brain structure and function and is differentially related to performance across specific cognitive domains.

Next, when assessing our and any other multimodal results, it is important to consider that that the parcellation of the brain impacts the relationship between brain structure and function and cognition (Dhamala et al., 2021; Messé, 2019). Here, we used a high-resolution brain atlas from Shen et al. (2013). High atlas resolution has been found to benefit investigations of the relationship between brain structure and function (Diez et al., 2015). Consequently, different model preference may be found using atlases with lower spatial resolution. In addition, previous work demonstrates that the topography of functional parcellation can contribute to the efficiency of its explanation of cognition (Kong et al., 2019). In the present work, we have used a functional atlas defined during resting state connectivity. Thus, it is likely that use of an atlas defined with cytoarchitectural information would also change the pattern of results. However, atlas use is a subject to wide debate as every atlas will focus on specific features of the brain (Eickhoff et al., 2018). For example, the atlas defined by Shen and colleagues (2013) is a whole brain atlas that jointly optimises subject and group level parcellation under constraint of local functional homogeneity. This means that the signal contained within each parcel is stable and less likely to include interfering signals across several parcels. In comparison, other research groups, like Bellec et al. (2010), have prioritised parcels that are stable at subject and group level, rather than locally homogenous parcels. Thus, it is clear that use of specific atlases will influence what information is prioritised by the models. The question is, however, why such information would change the patterns of results, and future work should carefully investigate how network characteristics differ across parcellations. For example, are specific atlases more sensitive to network modularity than others and does that sensitivity improve modelling of specific cognitive domains using structural, functional or joint connectivity.

## Conclusion

The present work has illustrated, for the first time, that the combination of information from structural and functional neural connectivity allows explanation of unique variation in performance in some, but not all, cognitive domains. The combination of structure and function produced a superior model of the neural bases of performance for the domains of Executive Function and Language, but not Self-regulation, Encoding, or Sequence Processing. This demonstrates that our use of combined structural-functional connectivity provides new insights concerning the different neural mechanisms involved in distinct cognitive domains. We have proposed that hierarchical cognitive organisation and interplay between direct and indirect functional connectivity is what drives the model preference in our research.

## Acknowledgements

This work was supported by the Biotechnology and Biological Sciences Research Council, UK (grant number: BB/M011208/1). Data were provided by the Human Connectome Project, WU-Minn Consortium (Principal Investigators: David Van Essen and Kamil Ugurbil; 1U54MH091657) funded by the 16 NIH Institutes and Centers that support the NIH Blueprint for Neuroscience Research; and by the McDonnell Center for Systems Neuroscience at Washington University.

