## Supplementary Materials for "Combination of structural and functional connectivity explains unique variation in specific domains of cognitive function"

**Supplementary material 1 - summary of behavioural measures**

| **Table 1. A summary of behavioural measures provided by HCP and utilised in this study. Measure name was followed by the name of the measure in the HCP dataset in italics.** | | |
| --- | --- | --- |
| **Measure Name** | **Subdomain** | **Description** |
| Picture Sequence Memory *(PicSeq_Unadj)* | Episodic Memory | Task requires encoding, storage and retrieval of non-verbal memory. Participant is presented with a sequence of pictures, pictures are scrambled and the participant is asked to reconstruct the order. |
| Dimensional Change Card Sort *(CardSort_Unadj)* | Cognitive Flexibility | Task requires planning and execution of behaviour with the goal of following constraints of rules. Participants is presented with picture cards following one rule (e.g. colour) then they are asked to reorder them following different rule (e.g. shape). |
| Flanker Inhibitory Control and Attention Task *(Flanker_Unadj)* | Inhibition | Task requires direction of attention in presence of conflicting cues presented in the environment. Participants is asked to indicate what direction the highlighted arrow is pointing in, while the surrounding arrows either point in the same direction (congruent condition) or opposite direction (incongruent condition) |
| Penn Progressive Matrices *(PMAT24_A_RTCR/(* *PMAT24_A_CR /24*100))* | Fluid Intelligence | Task requires solution of novel reasoning problems. Participants are presented with a sequence of matrices presenting a pattern and they are asked to choose a final square that will complete the pattern. |
| Oral Reading Recognition *(ReadEng_Unadj)* | Reading Decoding | Task requires decoding and auditory output of decoded words and letters. Participants are presented with a sequence of words of varying difficulty and they are asked to read them out loud. |
| Picture Vocabulary *(PicVocab_Unadj)* | Vocabulary Comprehension | Task requires decoding of written words and matching them with semantically closest picture. |
| Pattern Comparison Processing Speed *(ProcSpeed_Unadj)* | Processing Speed | Task assesses speed of information processing. Participants are asked to indicate as quickly ask possible if pictures match. |
| Delay Discounting *(DDisc_AUC_200 and*  *DDisc_AUC_40K)* | Self-regulation/Impulsivity | Task assess to delay of reward in order to benefit more from it. Participants are asked whether they would prefer to receive immediately half proportion of a monetary reward (either $200 or $40 000) or whether they would delay for a longer period (1 month to 10 years) to receive complete reward. |
| Variable Short Penn Line Orientation Test *(VSPLOT_CRTE / (VSPLOT_TC*  */24*100))* | Spatial Orientation | Task requires comparison of visuo-spatial information. Participants are asked to match orientation of blue line to the red line, while the location, orientation and length of the blue line varies. |
| Short Penn Continuous Performance Test *(SCPT_TPRT)* | Sustained Attention | Task requires extended dedication of attention to stimulus and matching of shapes. Participants are presented with lines that and the participants are asked to respond when the lines formed a letter or number. |
| Penn Word Memory Test *(IWRD_RTC/ (IWRD_TOT/40*100))* | Verbal Episodic Memory | Task requires encoding, storage and retrieval of verbal information. The participants are presented with a list of 20 words, then they are presented with another list containing original 20 words and 20 new words. The participants are asked if they had previously seen the word. |
| List Sorting *(ListSort_Unadj)* | Working Memory | Task requires encoding and storing of visual information for a short period of time in addition to retrieval and manipulation of information. The participant is presented with a series of picture and they are requested to sort them from memory in order of a feature (e.g. size). |

**Supplementary material 2 - Analysis of dense connectivity**

In section 2.2.1. and 2.2.2, a proportional 80% threshold was applied to SC and FC. In this part of supplementary material, the same analysis outlined in section 3.2. was applied to dense connectivity (without proportional threshold). In addition, the sign of negative FC correlations was preserved.

For the first time in the analysis, there was considerable AIC difference in favour of SC model relative to FC model of Sequence Processing (Sequence Processing ΔAIC = -4.1) Across all remaining cognitive domains there was considerable to strong preference in favour of FC relative to SC models (Executive function ΔAIC = 3.53, Self-regulation ΔAIC = 36.87, Language ΔAIC = 33.28, Encoding ΔAIC = 22.02). AIC differences of SC and FC models with respect to CC model for all cognitive domains are illustrated in Figure S4. Relative to SC, there was considerable preference for CC model of Sequence Processing, and strong preference for Executive Function, Self-regulation, Language and Encoding. Relative to FC, AIC values were strongly in favour of CC models of Executive Function, and Language and Sequence Processing. However, relative to CC model, AIC values strongly favoured FC model of Self-regulation. The AIC difference between CC and FC models of Encoding was barely worth a mention.

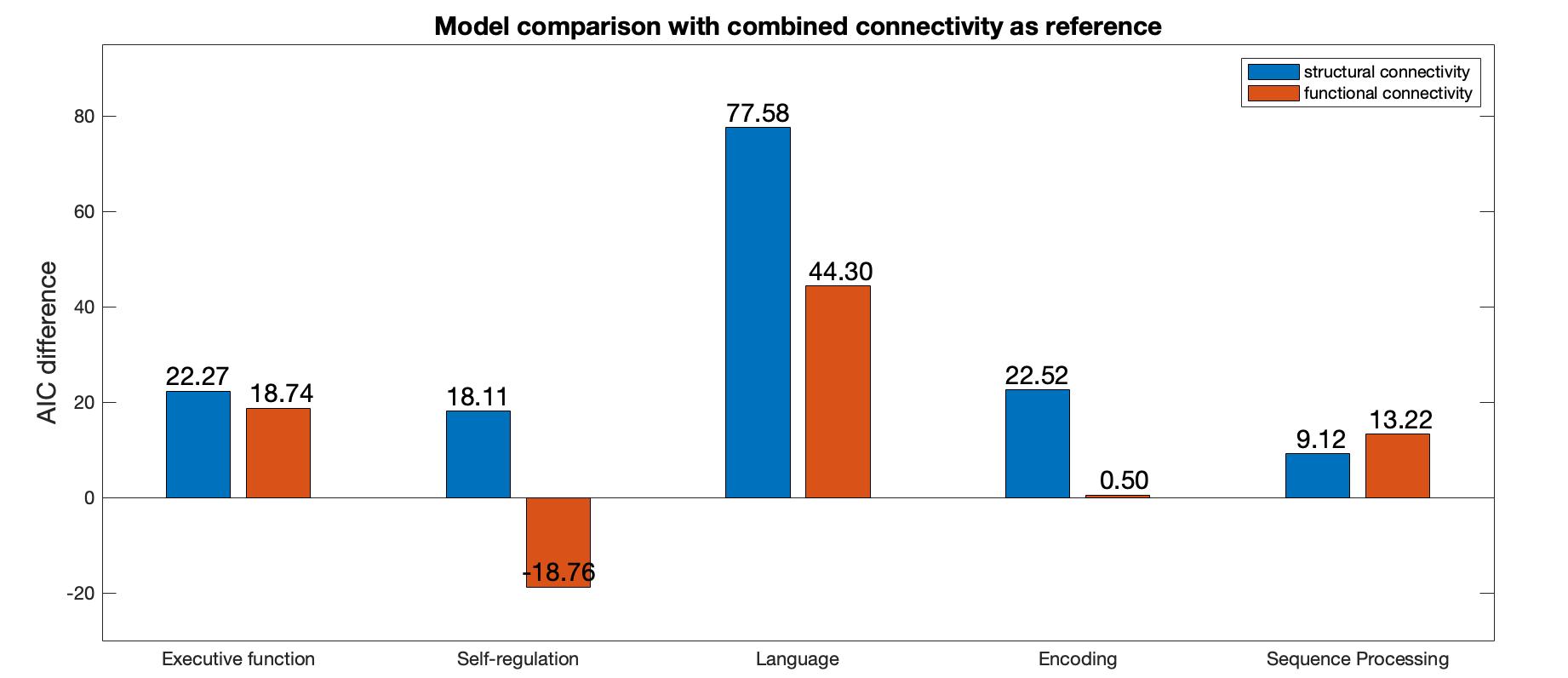

*Figure S4 AIC-based model comparison for each cognitive domain. The models were estimated from dense connectomes. The CC model was used as the reference model for model comparison. Bars with positive values indicate preference for the reference model.*

Figure S5 illustrates the explained variance of the models under consideration. Across domains CC models achieved weak to moderate accuracy in prediction of cognitive data (Ordinary R-Squared range 0.05-0.62). The FC model explained most variance in Self-regulation and Encoding, whereas Executive Function, Language and Sequence Processing were best explained by the CC model.

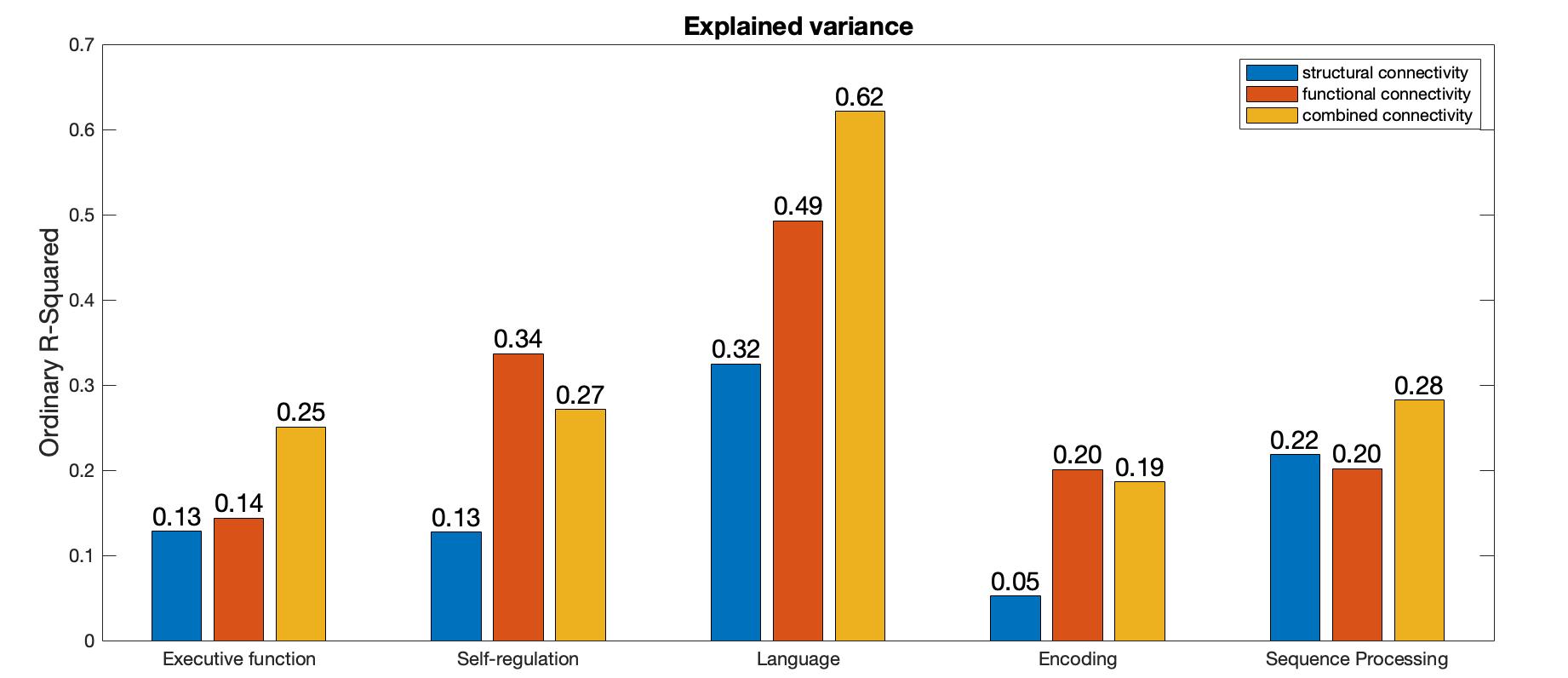

*Figure S5 Bar graphs illustrate the variance explained by dense SC, FC and CC models of each cognitive domain.*

Figure S6 illustrates the prediction error generated for unseen sample of 49 participants. FC had generated least prediction error for models of Executive Function, Language. It produced approximately the same amount of error as CC models of Sequence Processing. CC produced least out of sample prediction error of Self-regulation.

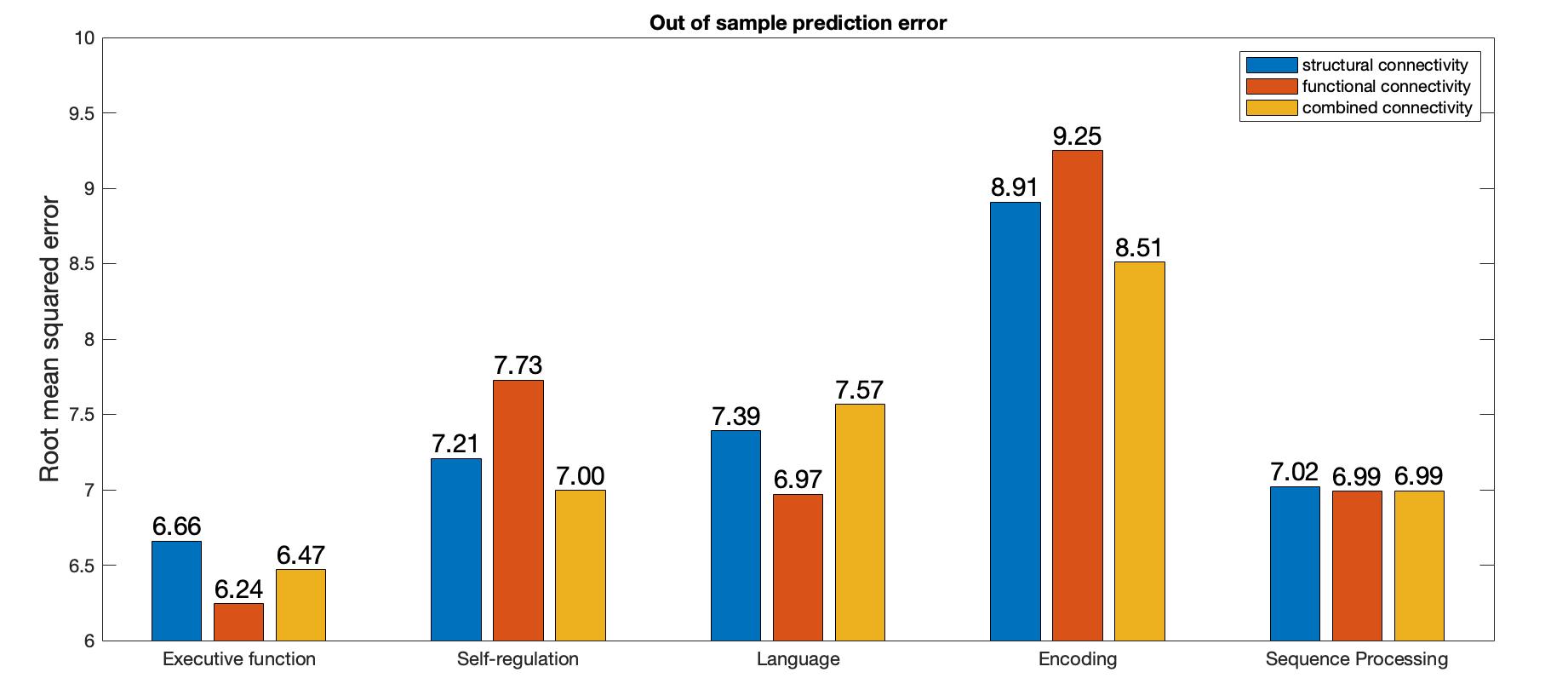

*Figure S6 Out of sample (N = 49) prediction error generated by dense connectivity models, as measured by root mean squared error.*

**Supplementary material 2 – Analysis of FC preserving sign of negative correlations**

In section 2.2.2., negative correlations of FC matrices were transformed to positive by taking their absolute values, then a proportional 80% FC threshold was applied. This supplementary material presents the results of analysis outlined in section 3.2. when the same threshold was applied but the sign of connectivity was restored after thresholding.

Across all cognitive domains there was considerable to strong preference in favour of FC relative to SC models (Executive function ΔAIC = 8.86, Self-regulation ΔAIC = 31.96, Language ΔAIC = 28.55, Encoding ΔAIC = 24.67, Sequence Processing ΔAIC = 42.04). AIC differences of SC and FC models with respect to CC model for all cognitive domains are illustrated in Figure S1. Relative to SC, there was weak preference for CC model of Encoding, considerable preference for Executive Function and strong preference for Self-regulation, Language and Sequence Processing. Relative to FC, AIC values were considerably in favour of CC models of Self-regulation and Language. However, relative to CC model, AIC values strongly favoured FC models of Encoding and Sequence Processing. The AIC difference between CC and FC models of Executive Function was barely worth a mention.

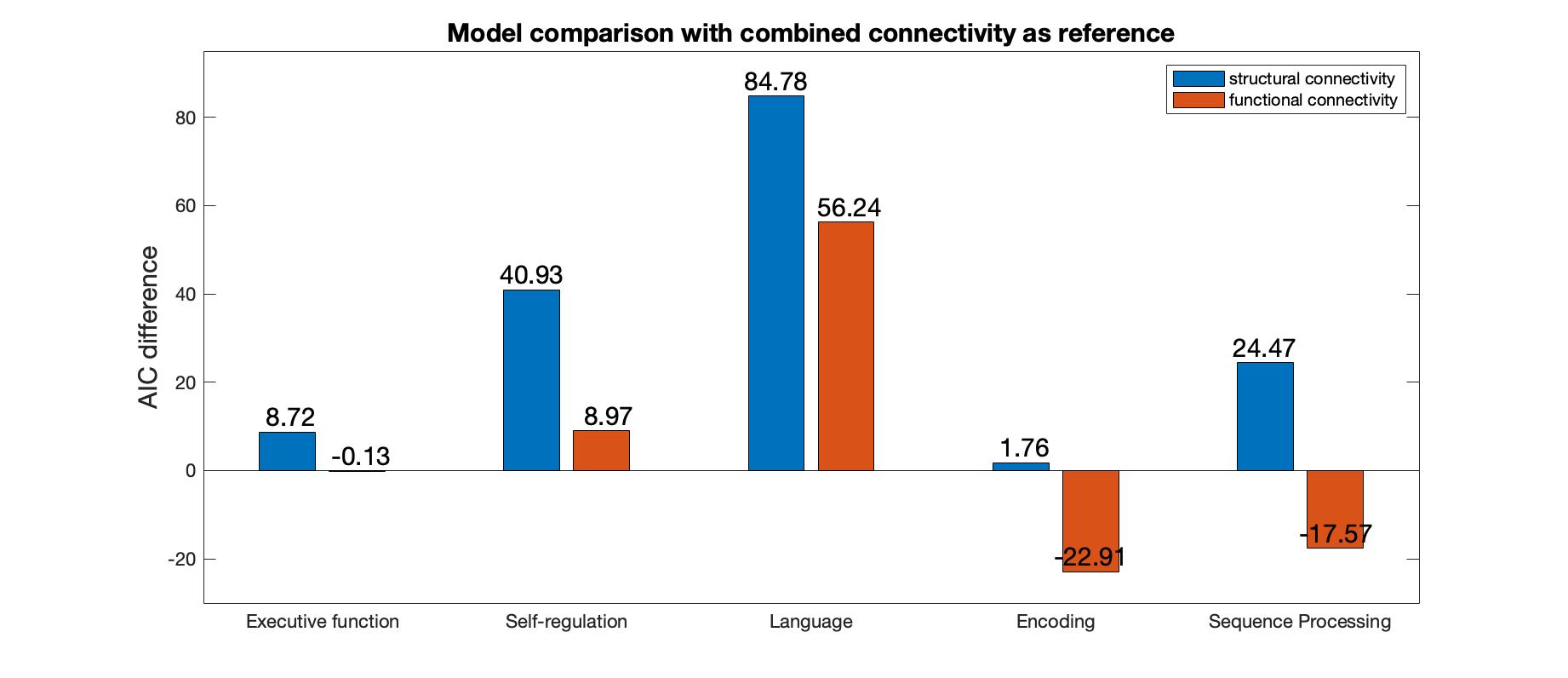

*Figure S1 AIC-based model comparison for each cognitive domain. The FC was thresholded and signs of negative correlations were restored. The CC model was used as the reference model for model comparison. Bars with positive values indicate preference for the reference model.*

Figure S2 illustrates the explained variance of the models under consideration. Across domains CC models achieved weak to moderate accuracy in prediction of cognitive data (Ordinary R-Squared range 0.10-0.66). The FC model explained most variance in Encoding and Sequence Processing, whereas Executive Function, Self-regulation and Language was best explained by the CC model.

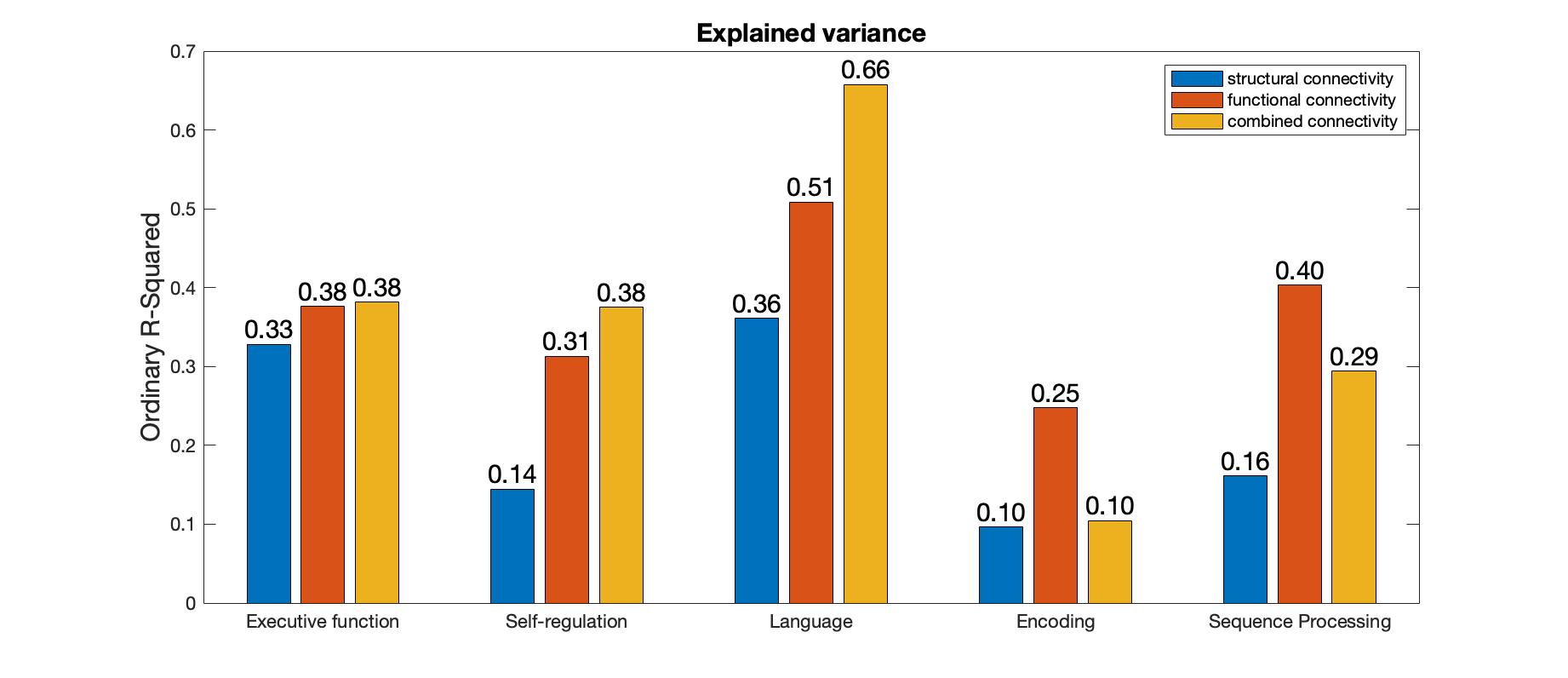

*Figure S2 Bar graphs illustrate the variance explained by SC, FC and CC models of each cognitive domain. The FC was thresholded and signs of negative correlations were restored.*

Figure S3 illustrates the prediction error generated for unseen sample of 49 participants. As illustrated in Figure 6, SC had highest generalisation skills for Language, FC performed best for Executive Function, Self-regulation and Sequence Processing, CC performed best for Encoding.

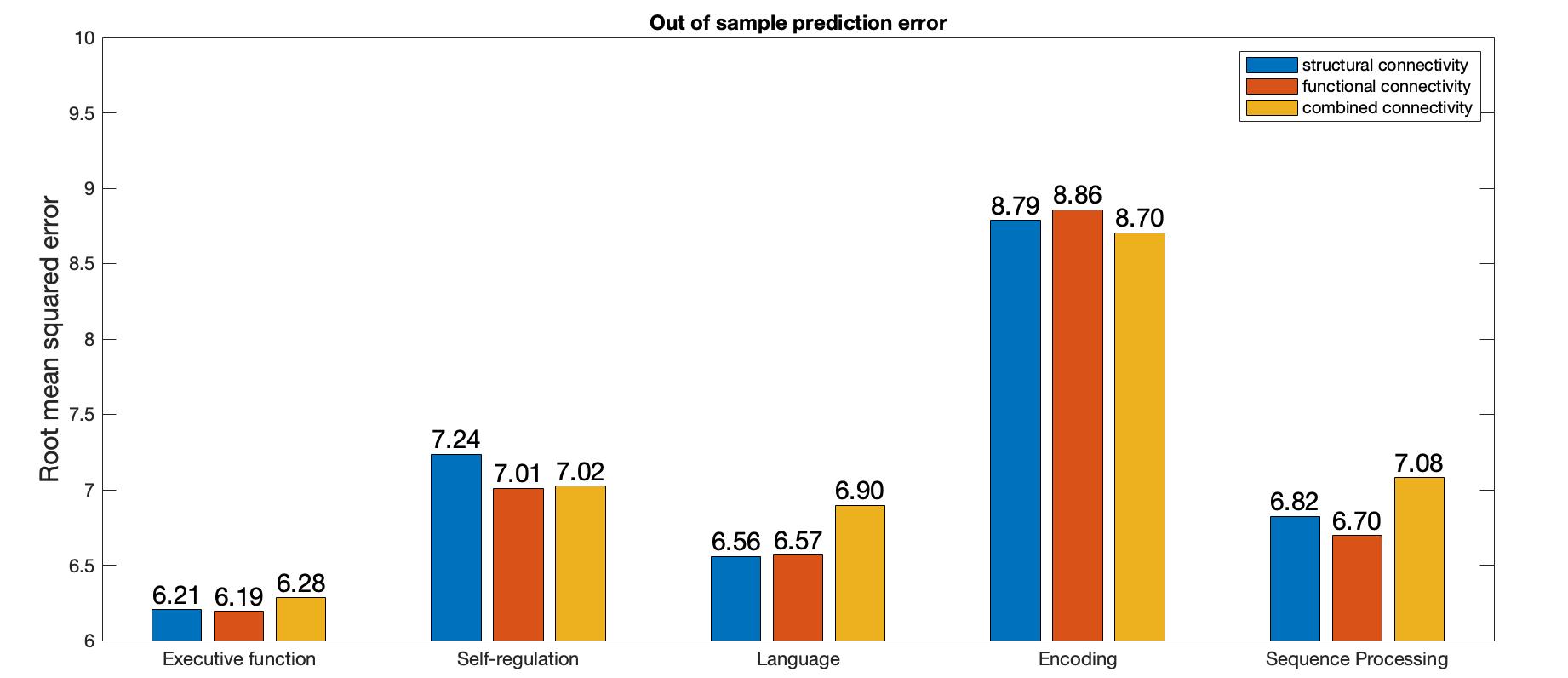

*Figure S3 Out of sample (N = 49) prediction error, as measured by root mean squared error. The FC was thresholded and signs of negative correlations were restored.*
